# Creating ethnicity-specific reference intervals for lab tests from EHR data

**DOI:** 10.1101/213892

**Authors:** Nadav Rappoport, Hyojung Paik, Boris Oskotsky, Ruth Tor, Elad Ziv, Noah Zaitlen, Atul J. Butte

**Affiliations:** Institute for Computational Health Sciences, University of California, San Francisco, CA 94143 US; Department of Pediatrics, University of California, San Francisco, San Francisco, CA 94143 US; Korea Institute of Science and Technology Information, Biomedical Convergence Technology Research Division, Biomedical HPC Technology Research Center, Deajeon 34141, South Korea; Rabin Medical Center, Beilinson Hospital, Petach Tikva, Israel 4941492; Institute for Human Genetics, University of California, San Francisco, San Francisco, CA 94143 US

## Abstract

The results of clinical lab tests are an essential component of medical decision-making. To guide interpretation, test results are returned with reference intervals defined by the range in which 95% of values occur in healthy individuals. Clinical laboratories often set their own reference intervals to accommodate local population and instruments variations. This approach is costly and can be biased. We describe a novel data-driven method for using electronic health record data to extract healthy patients’ information to define reference intervals. We found that the distributions of many clinical lab tests differ among self-identified racial and ethnic groups (SIREs) in healthy patients. Finally, we derived SIRE-specific reference intervals and provide evidence that these intervals have clinical prognostic value. Specifically, we show that for two lab tests, serum creatinine level and hemoglobin A1C, SIRE-specific reference intervals are more predictive for need for dialysis and development type 2 diabetes than existing reference intervals.

**One Sentence Summary:** A novel method for defining population-specific reference intervals of common clinical laboratory tests from electronical health records has better prognostic value than existing reference intervals.

## Introduction

Clinical laboratory tests contribute to medical diagnoses and interventional decisions. They also provide insight into physiological states that are not directly observable. However, differences in population demographics, geography, and lab instruments can alter distributions of test results (*1, 2*). Each clinical laboratory is expected to define reference intervals for each lab test (*1, 3*). Many hospitals, including UCSF Medical Center, base these intervals on published specifications and/or data from healthy individuals from the communities around the hospital. The intervals are incorporated into the EHR system and used to define normal/abnormal results. Reference intervals are typically defined as the interval where 95% of test results in healthy individuals occur (*4*). The current gold-standard approach is to collect 120 healthy samples to estimate reference intervals (*3, 5*). However, reference samples are not always be possible to collect, as a diverse set of healthy individuals may not volunteer for unnecessary clinical procedures (*6*). There is also no general agreement on how to define healthy individuals, and labs often use samples that are “easily collected,” such as college students in academic clinics or internal laboratory staff. Some laboratories do not collect their own samples, but rather rely on reference intervals from the literature or equipment manufacturers (*3*).

Distributions of normal lab test values can vary between self-identified races and ethnicities (SIRE). To account for these differences, some test results use SIRE-specific reference intervals (*7*). For example, serum creatinine has different distributions for African-Americans and Europeans (*8-10*). Estimated glomerular filtration rate (eGFR), a widely-used test of kidney function, is a calculation based on four variables: serum creatinine, age, sex, and race (*11*). However, in this test, race is binary (African-American or European), and recent work suggests that more granular categories and/or genetic ancestry could improve eGFR scaling (*12*). In other instances, differences in test distributions between SIRE are known in theory, but reference intervals are not altered in practice (*13, 14*). For example, a genetic variant in the Duffy Antigen Receptor (*15*), common in African-Americans but rare in European-Americans, induces a one standard deviation drop in mean neutrophil count. It is the basis of *benign ethnic neutropenia* (*16*), but neutrophil counts are reported with the same reference interval for all SIREs.

This work considers changes to the definitions of normal/abnormal test results when reference intervals come from (1) existing reference intervals, (2) a complete collection of healthy individuals, or (3) a population of matching individuals. We have developed a method for defining reference intervals using data taken directly from the electronic health record (EHR) system at the University of California, San Francisco (UCSF) medical center. This method reduces the need to collect samples from reference populations, matches the reference to local patient demographics, and incorporates instrument noise. EHR-based research may have advantages over traditional clinical studies (*17*). This is because EHR-based studies use data acquired under routine conditions, whereas large studies use measurements acquired by following research protocols. Furthermore, EHR-based reference interval calibration permits direct examination and accommodation of test distribution differences between SIREs. We applied our approach to the 50 most commonly ordered lab tests in the UCSF patient population and found that 50% differed between SIREs. Most importantly, we found that altering SIRE-specific reference intervals for serum creatinine and hemoglobin A1C (HbA1c), improved the predictive power for eventual diagnoses of kidney disease requiring dialysis and diabetes mellitus (type 2 diabetes; T2D) over existing reference intervals.

## Results

We examined EHR data from ~920,000 patients (26 million encounters and 81 million results) seen between August 2012 and September 2017. Table 1 shows basic demographic information for this population. We selected our main cohort from the EHR based on inclusion and exclusion criteria, as follows. To capture healthy patients, we selected outpatients with a diagnostic International Classification of Diseases, Tenth Revision, Clinical Modification (ICD10) code from a predefined list (Supplementary Table S1) and excluded any encounter with ICD10 codes not on the list. Out of 23 million encounters, there were 13,559 in 10,950 healthy adults (18 – 60 years old). A total of 179,774 lab test results were recorded (Table 2). We ensured that lab instruments were not changed during the collection period.

**Table 1.** Summary statistics of patients in UCSF EHR database.

|  | **Median** | **Mean** | **Standard deviation** |
| --- | --- | --- | --- |
| **Age** | 42 | 41.66 | 23.98 |
| **Encounters per Patient** | 7 | 27.19 | 61.48 |
|  | **Female** | **Male** | **Unknown/ Unspecified** |
| **Sex** | 505400 | 415888 | 1308 |

| **Race \ Ethnicity** | **Hispanic or Latino** | **Not Hispanic or Latino** | **Unknown/Declined** |
| --- | --- | --- | --- |
| **Native American or Alaska Native** | 562 | 2337 | 193 |
| **Asian** | 889 | 88912<sup>‡</sup> | 3510 |
| **Black or African American** | 937 | 39805<sup>‡</sup> | 2309 |
| **Native Hawaiian or Other Pacific Islander** | 295 | 12527<sup>‡</sup> | 1095 |
| **Other** | 70814<sup>‡</sup> | 47184 | 7927 |
| **Unknown/Declined** | 8080 | 15912 | 190276 |
| **White or Caucasian** | 20513<sup>‡</sup> | 351787<sup>‡</sup> | 20150 |

<sup>‡</sup> Indicates major race-ethnicity group used in the current analysis.

**Table 2.** Patients, Encounters, and Measurements.

|  | Healthy cohort<sup>a</sup> | General cohort |
| --- | --- | --- |
| **Ages** | 18 – 60 | 18 – 72 |
| **Number of patients** | 11,495 | 629,056 |
| **Number of encounters** | 14,387 | 19,372,922 |
| **Number of lab tests** | 189,907 | 58,593,901 |

<sup>a</sup>Based on ICD10 diagnosis codes listed in Table S1.

### Labs reference intervals can be computed from EHR data

We collected data from the 50 most common lab tests having at least 120 healthy samples for at least one sex from the UCSF EHR. Outlier values defined as values exceeding the first or the third quartile by 3 times the center inter-quartile range were considered outliers and removed (see Methods). Reference intervals were then defined as the central 95% of values.

EHR-calibrated reference intervals defined by our method did not substantially differ from existing ones (Supplementary Table S2). For example, UCSF’s current reference interval for creatinine in females aged ≥19 years is 0.44 – 1.0 mg/dL. Based on 2,974 healthy females aged 19-60, we defined a new reference interval as 0.48 – 0.97 mg/dL (Supplementary Table S2). Our interval would reclassify 19,837 (3.2%) previously normal measurements to below the lower threshold, and 9,450 (1.5%) measurements to above the higher threshold. Similarly, the existing reference interval for white blood cell counts (WBC) for females aged ≥21 years is 3.4 – 10 x10^9^ cells/L and our EHR-calibrated reference interval (3,532 subjects) was 3.3 – 10.5 x10^9^ cells/L. Here, 4,010 measurements (0.6%) would reclassify from low to normal, and 18,983 (2.8%) from high to normal (Supplementary Table S2). The average absolute change in intervals was 21%, with a standard deviation of 21% (see Methods). These changes would affect 6.1% of all measurements: 4.0% of out-of-range measurements will be reclassified as normal and 2.1% of normal measurements would exceed the new thresholds.

By comparing the ratios of the new intervals and the original ones vs. sample size, we found that the intervals of two lab tests had a greater than two-fold change (Figure S1). The existing reference range for direct bilirubin is ≤ 0.3 mg/dL for females and males. We identified a range of ≤0.1 mg/dL for both females and males independently, which would be a 3-fold narrowing of the reference interval. The existing reference range for C-reactive protein (CRP) is ≤6.3 mg/L for both sexes in UCSF’s reference interval, while our EHR-calibrated method gave upper threshold/95th percentile values of 15.52 and 13.0 mg/dL for females and males, implying a two-fold increase over the UCSF reference intervals. CRP is very sensitive to infection. Part of our healthy cohort are patients who showed up for immunization which may affect CRP levels. We recalculate an EHR-calibrated reference interval without immunization patients, and found upper threshold for females (males) to be 17.28 (13.87) mg/L based on 162 (118) healthy samples.

### Lab test distributions differ across populations

General practice for defining reference intervals requires separate categories for males and females and for age groups when appropriate. Few lab tests have reference intervals for races and ethnic groups currently in use in the clinic (e.g., creatinine). We had male-specific data from 46 tests and female-specific data from 42 tests that also had at least two SIREs with ≥50 healthy individuals. For each test, we compared the distribution of healthy measurements from different SIREs adjusting for age and stratifying by sex and tested for differences using ANOVA (see Methods). Table 3 shows the top 10 results. The complete set of results, listed in Supplementary Table S3, shows that many lab tests differed between SIREs. Out of 88 lab tests, 45 (51%) were significantly different in different SIREs (ANOVA p-value < 0.05 after Benjamini-Hochberg correction for multiple testing). The nonparametric Kruskal-Wallis test also found statistically significant differences between SIREs in 68 (77%) lab tests (Supplementary Table S3). We concluded that many of tests have different distributions across SIREs.

**Table 3.**
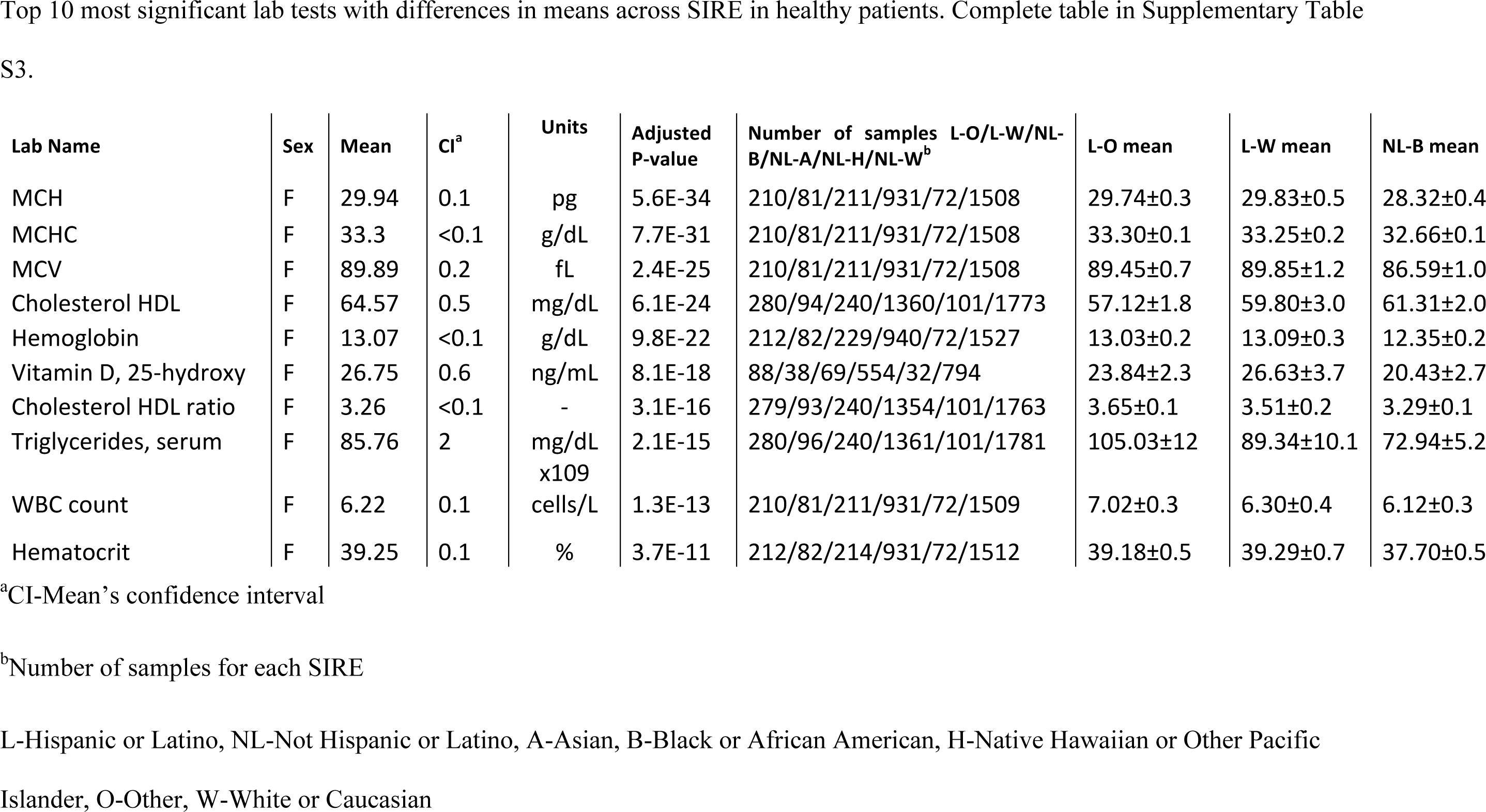
Lab tests and adjusted ANOVA p-values.

### Distribution of Clinical Measurements within Healthy Individuals

Results for several lab tests are known to have distribution differences between SIREs. Serum creatinine is an example (*8*). We evaluated this test in detail. Healthy non-Hispanic African-Americans had the highest means, followed by non-Hispanic Europeans and Hispanic Europeans, with non-Hispanic Asians having the lowest means (Figure 1A), consistent with the previous publications(*7, 8*). The difference in creatinine levels between female SIRE was significant (ANOVA p-value = 4.1×10^−47^, KW p-value = 1.7×10^−10^, Supplementary table S3), and remained significant after excluding non-Hispanic African-American samples (p-value = 2.7×10^−6^). The difference in males was not significate according to ANOVA test (p-value = 0.24), but was according to KW test (p-value = 8.6×10^−49^)

**Fig. 1.**
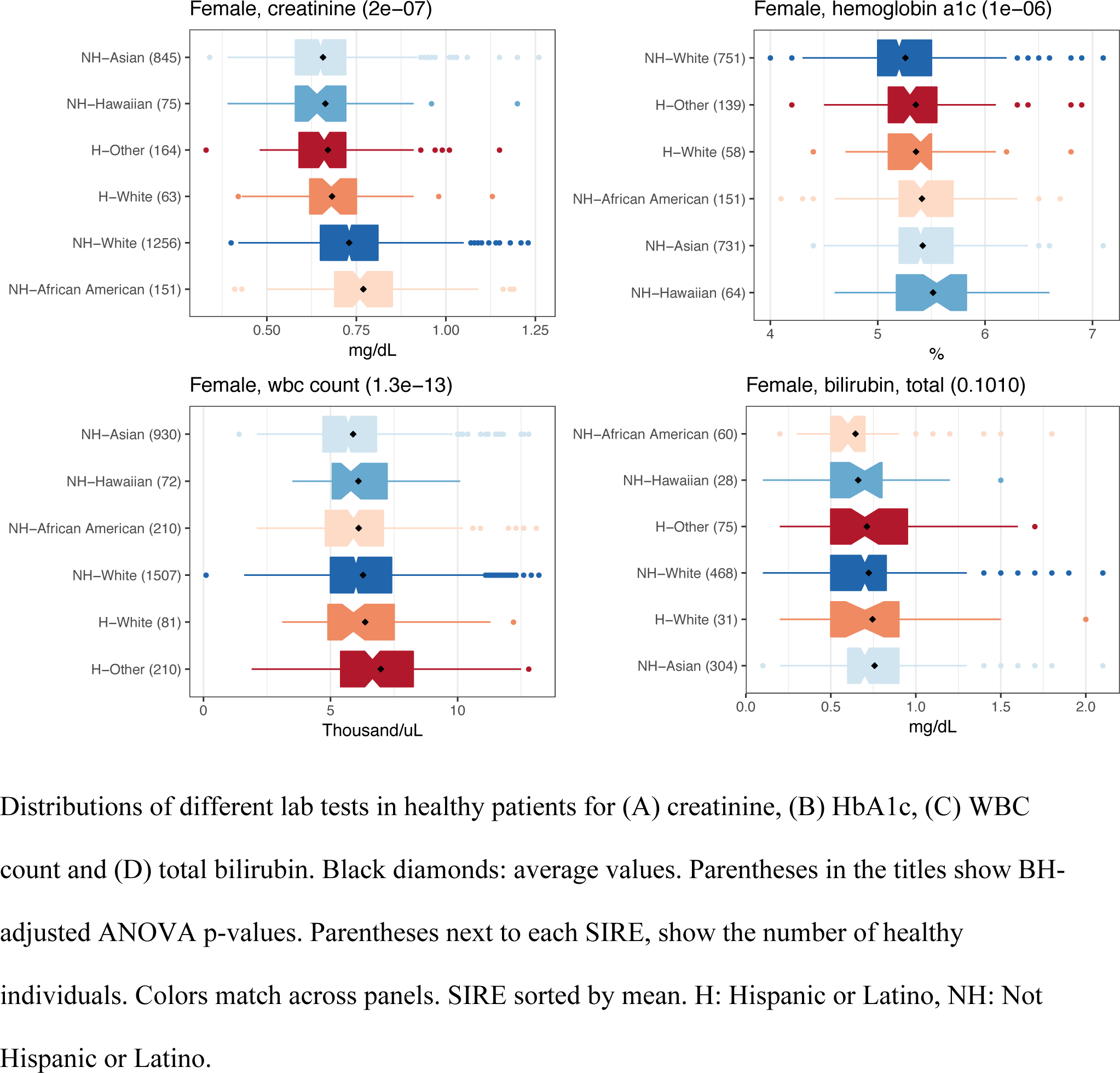
Difference in lab test measurements among healthy patients in different SIRE groups.

Other studies have also suggested differences in total bilirubin levels between SIREs (*18*). We tested this possibility and found no difference (ANOVA p-value 0.1 and 0.25 for females and males respectively; Figure 1D and Supplementary Table S3). We also found that healthy African-Americans had a lower average WBC count than healthy Europeans (T-test p-value = 0.045), as has been shown (*15, 19, 20*). We also found a lower WBC count in non-Hispanic Asians compared to Europeans and Hispanics (T-test p-value = 1.3×10^−11^, Figure 1 and Supplementary Table S3). HbA1c levels in our data were higher in African-Americans compared to Europeans in accordance with previous work (*21, 22*), but a recent study found the opposite: African-American had lower values of HbA1c (*23*). To the best of our knowledge, the majority of these differences are novel findings and are not in use in the clinic.

### Evaluation of Adjusted Reference intervals

Previous SIRE-specific reference intervals were adopted on the basis of distribution surveys (*8, 24, 25*). We determined whether using different reference intervals improves the predictive power or the prognostic value of lab tests. First, we modeled the relationship between future health outcomes and test results in the entire adult patient cohort (healthy/non-healthy; aged 18 to 72 years; Table 2). We selected three lab tests with well-known outcomes of abnormal results (HbA1c, serum creatinine and WBC). HbA1c provides information about average blood glucose levels over the past 3 months and is associated with T2D (*26*). A 1%-unit increase in HbA1c levels doubles the risk for T2D (*22*). Serum creatinine measures kidney function and elevated levels of this molecule indicate kidney disease (*27, 28*). WBC are elevated in reaction to bacterial infection (*29, 30*). Classification of first available measurement of each patient as low/normal/high according to the current reference range was associated with a specific outcome (log-rank test (*31*) p-value < 1×10^−16^; Figure 2). To test whether different SIRE groups had different outcomes in spite of the same test results, we compared 4 nested Cox regression models where the dependent variable was the outcome (dialysis for creatinine, T2D for HbA1c and infection for WBC) and the independent variables were (i) lab values and sex (current practice); (ii) lab values, sex and age; (iii) lab values, age, sex, and SIRE; and (iv) an interaction term between SIRE and lab test value in addition to (iii). An interaction term in a regression model enables a separate relationship between lab test and outcome for each SIRE and if significant, suggests that SIRE-specific intervals have additional predictive power. We found that adding an interaction term between SIRE and the lab values increased the predictive performance (likelihood ratio test’s p-values = 5.26×10^−110^, 1.4×10^−221^ and 8.9×10^−9^ for creatinine, HbA1c and WBC respectively; Table 4). Here again, we confirmed that significant difference in creatinine levels was not derived solely by non-hispanic African-Americans’ higher mean by rerun the test after eliminating these samples (likelihood ratio test p-value = 4.18×10^−99^). Because HbA1c ≥6.5% is generally considered as diagnostic of T2D (*32*), we repeated the test after removing patients whose first HbA1c result was ≥6.5%, and reconfirmed the significance (likelihood ratio test’s p-value = 9×10^−9^).

**Table 4.**
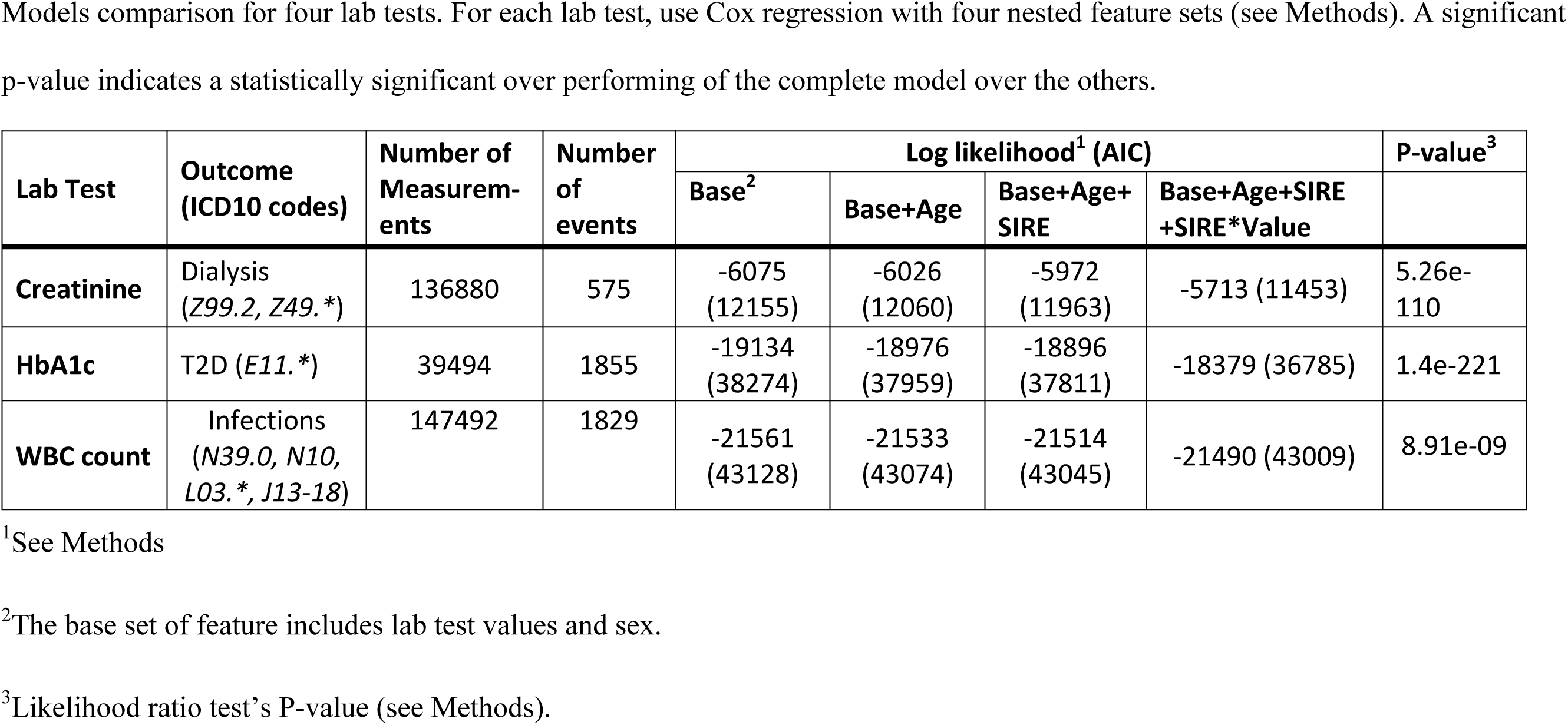
Comparison of specific outcomes regressions

**Fig. 2.**
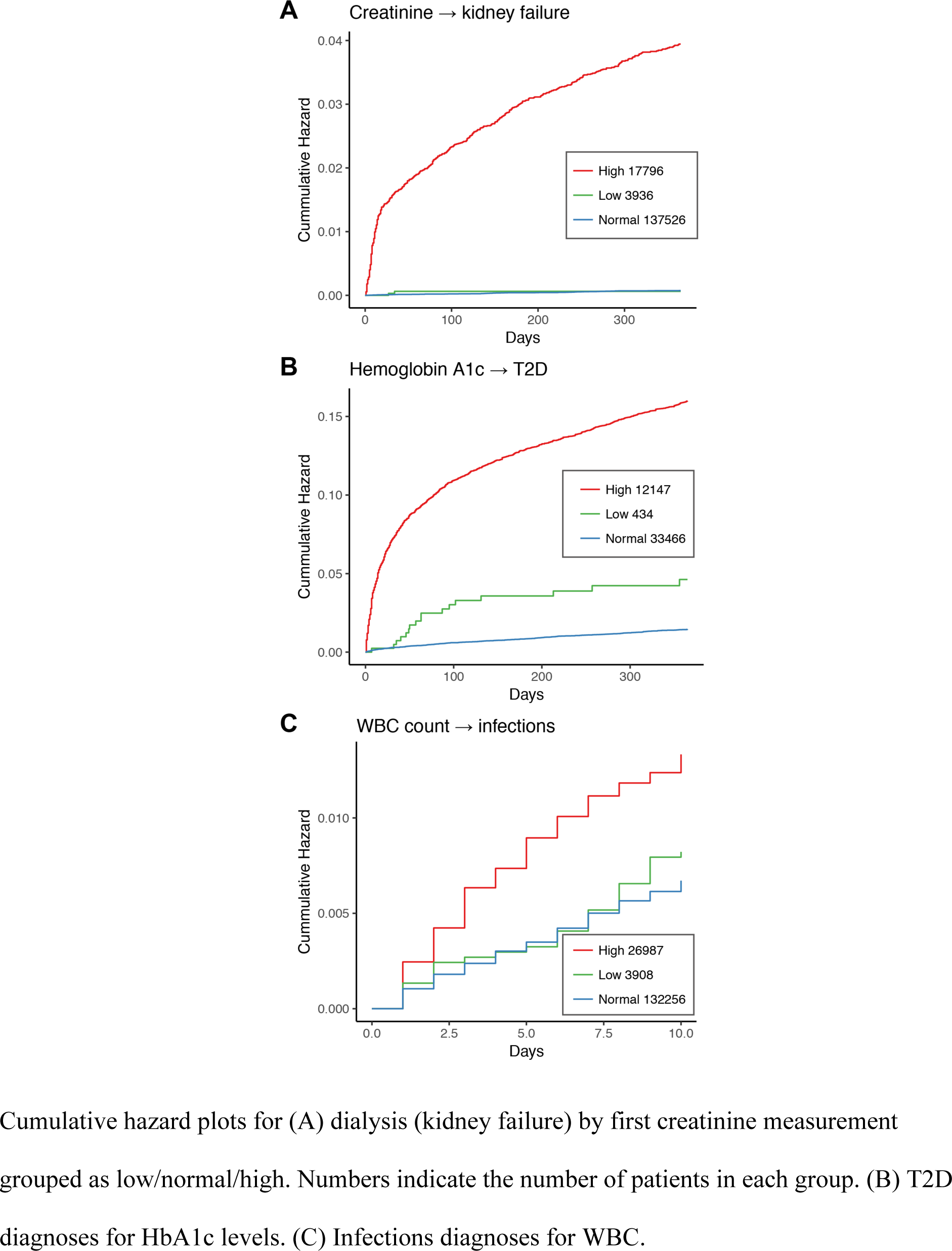
Cumulative hazard rates.

In a similar approach using logistic regression instead of Cox regression, we counted the number of patients who experienced the outcome of interest within a year of positive test results (positive patients) and those who did not (negative patients). Using the interaction term improved positive predictive value by 2% (0.35, 0.68 for creatinine and HbA1c respectively; Supplementary Table S4). Here, the predictive power of WBC found to be low. We conjected that this is due to the broad health conditions that may lead to elevation in WBC. We concluded that a SIRE-specific reference interval correlates with outcomes better than common practice, which does not include SIREs. It was also better than adding SIRE as a factor without accounting for interactions between SIREs and lab tests.

### Reference intervals should be stratified according to population demographics

We next examined the effect sizes of SIRE-specific reference intervals. SIRE- and sex-specific reference intervals were refined. For each lab test, sex and SIRE, reference interval was defined as central 95% of values of matching healthy patients. The number of measurements in the low/normal/high group for each reference intervals considered was calculated for each SIRE and lab test. Among SIREs with at least 120 samples, 935,900 measurements (4.03%) originally considered abnormal were reclassified as normal, and 598,356 (2.57%) originally normal measurements were abnormal under SIRE-specific reference intervals (Supplementary Table S5).

For example, 19,593 non-Hispanic Europeans and 3,450 non-Hispanic African-Americans had HbA1c measurements. Using the first measurement per subject, 4,309 (22%) Europeans and 1372 (40%) African-Americans exceeded 5.6%, which is the upper threshold of the existing reference interval. Alternatively, only 955 (4.9%) non-Hispanic Europeans and 330 (9.6%) non-Hispanic African Americans exceeded it using an EHR-calibrated threshold of 6.2% (Supplementary Table S2). When a SIRE-specific threshold (Supplementary Table S5) was used 1451 (7.4%) and 256 (6%) of non-Hispanic Europeans and non-Hispanic African-Americans had values above the threshold. These findings demonstrate that a substantial number of individuals are newly categorized to abnormal from normal or vice versa when SIRE-specific reference intervals are constructed.

### Population-specific intervals have clinical effects

The findings above do not mean that altering reference intervals will also affect predictive power, and we considered this point. For a given lab test, we compared three simplified models. We used two binary variables representing three states: lab test measurement 1) below the lower threshold, 2) within the reference interval or 3) above the upper threshold. The thresholds were defined as follows: (i) the original, in use at UCSF; (ii) our EHR-calibrated interval based on healthy encounters; and (iii) an EHR-calibrated interval per SIRE. We used Cox regression and compared performance based on Akaike’s information criterion (AIC; see Methods) (*33*).

Dialysis prediction from creatinine abnormalities was improved when our reference interval was used compared to the one currently in use at UCSF (AIC reduced from 8,713 ±12 to 8,600 ±12) and slightly further improved when the SIRE-specific range was used (AIC: 8,574 ±12). Similarly, the predictive power of HbA1c abnormalities to progression to T2D improved when our reference interval was used (AIC reduced from 27,330 ±20 in the UCSF interval to 25,815 ±20), but was slightly worse when SIRE-specific range was used (AIC=25,972 ±20). Infections prediction were not improved by our reference interval compared to the one currently in use (AIC: 32,805 ±23 to 32,802 ±23), but was improved when SIRE-specific range was used (AIC: 32,774 ±23). This finding suggests that alternatives reference intervals may improve prognostic value for some SIRE-specific distributions.

## Discussion

We have created a novel approach to defining reference intervals from existing data. We found that distributions of many lab test results differed among SIREs and that accounting for SIRE improved the prognostic value of some tests.

EHR-calibrated intervals may provide more accurate estimates of “real” normal values because they are based on a larger number of samples than the 120 required by current gold-standard methods. They also avoid active collection of data. However, their use may lead to inclusion of abnormal values that are misclassified as healthy. For some lab tests, reference intervals may not be defined as the inner 95% of measurements (e.g., national society guidelines); therefore, EHR-calibrated ranges may differ substantially. EHR-calibrated reference intervals should be carefully implemented and attention must be given to subject clinical conditions that may affect tests results.

### SIRE-specific differences

Some of the environmental or genetic factors that may affect lab test results are known (*15, 34*). For example, the variant rs2814778 causes benign ethnic neutropenia in African-Americans. However, our finding that WBC levels were lower in normal non-Hispanic Asians cannot be explained by this variant, as its frequency is 0.82 in African-Americans and <0.001 in Asians as well as in Europeans (*35*).

The upper threshold of EHR-calibrated CRP was double that of the UCSF reference interval. The existing CRP range was defined in 1972 based on 145 samples (*36*). A study from 2005 (*37*) based on 8,874 samples from NHANES data (*9*) found that 10% of adult samples have CRP values ≥10.0mg/L. This supports our finding that a threshold of 6.3mg/L does not cover 95% of normal samples, and indicates that revising the current reference interval may be in order.

Our results suggest that altering reference intervals to make them more accurate may influence clinical prognosis. CRP is very sensitive to any illness, and its levels rise rapidly within hours of infection (*38-41*). Our subjects might have been exposed to infection, which may have affected the distribution of the results, and this may be true in the other studies as well.

Different sub-populations may seek care for different diseases due to genetic and environmental factors. This fact may drive differences in the distributions of test results, even if the healthy interval of these distributions is identical across populations. Although we found that >50% of tests varied by SIRE, only one of these, creatinine, is adjusted for SIRE in practice. Even in this case, we found that the simplified version of African American vs. non-African American misses heterogeneity, particularly the lower levels of creatinine among Asian Americans. African-Americans’ risk for mortality from stroke is two-fold greater than that of Europeans (*42*) and the prevalence of hypertension is increased in this group (*43*). In this case, setting a different reference interval for African-American patients may be the wrong approach.

Our findings suggest use of reference intervals based on healthy baselines determined for each group separately. However, it is not necessarily the case that a group-specific reference interval conveys more diagnostic information, even if the healthy group-specific distribution is different from healthy distributions in other populations. For example, healthy individuals from one population may have higher LDL cholesterol due to genetic and environmental factors such as diet. However, this does not necessarily mean that the risk of a cardiovascular event at a given level of LDL cholesterol differs between SIREs (*44-46*). Furthermore, the same level of an entity in different SIREs can convey different risks for specific outcomes. For example, the risk of developing diabetes and cardiovascular disease in African-American women with upper body obesity (UBO) is lower than European women with an equivalent degree of UBO (*47*).

Having a population-specific interval is just one step toward precision medicine, where an individual is diagnosed and treated not only compared to the general population but to a narrower population that represents that individual and his or her background. Even if SIRE-specific reference intervals are used, it is possible that reference intervals should be scaled based on the patient’s specific context. For example, SIRE-specific ranges for WBC may more accurately identify current infection, but may not be useful when determining when to stop chemotherapy. To get a more precise reference interval one should consider intra-individual and inter-individual variations in the test (*48*), but this type of information is not always available, as healthy patients tend not to come to the clinic, and the physician does not order tests for such patients unless part of preventive care.

Many clinical researchers need a background cohort that serves as a control and as a statistical background. The control cohort should match the intervention cohort as closely as possible except for the treatment condition, and collecting control data can costs as much as case data. We have shown a novel way to define and extract background cohorts from an EHR system which might be useful for other case-control studies.

### Limitations of the study

While there are advantages to using a data-driven method to determine reference intervals, there are also limitations to our approach. For example, patient numbers may insufficient for less commonly ordered tests and for tests that are usually not ordered for healthy patients. An additional limitation is using EHR diagnosis codes to choose healthy patients. It is possible that diagnoses were missing for some of our patients. Some subjects might have been taking medications or had an illness that was not recorded during the encounter when tests were ordered. Our healthy cohort may also have included undiagnosed diabetic or pre-diabetic patients who did not met our exclusion criteria, which would falsely increase our HbA1c reference interval. In addition, ICD10 is also used for billing and reimbursement purposes, undercoding, or assigning fewer ICD10 codes or codes with lower costs, is common among physicians to avoid audits or minimize costs (*49, 50*). Despite these limitations, our method does at least as well the standard method for sample collection which lacks a standard for defining healthy volunteers (*51, 52*). In addition, some lab test results change dramatically in different normal conditions. For example, calcium and phosphate levels have a circadian rhythm (*53-55*), creatinine is influenced by hydration (*56*) and glucose by food intake. None of these conditions were controlled here, but aren’t considered in the current scheme of determining reference intervals either.

We grouped patients based on self-identified race and ethnicity, which may not be accurate. First, false reporting is common (*57, 58*). Also, some groups, such as Latinos and African Americans, have mixed heritage, and varying genetic ancestry in these individuals may lead to inaccurate assessments if they are treated as one group (*12*). In addition, many patients identify with two or more racial/ethnic categories. An extension of the current work will define reference ranges for more population categories (or even a continuous space of patients), and may also include patient genome information. However, environmental factors such as diet may also affect lab test results (*59*).

Reference intervals are a statistical view of a population. Alternatively, decision limits are obtained from epidemiological studies, and are used to determine the likelihood that a given disease is present or in higher risk. In contrast to reference intervals, they cannot be extracted from a healthy local population. If defined and applicable for a given situation, they should be used instead of a reference interval. For example, the National Cholesterol Education Program states that >15% of young adult men have a cholesterol HDL <35 mg/dL. However, the desired value is ≥40 (*60*) in order to decrease the risk for coronary heart disease.

A point to be considered on the path to precision medicine is that some modifications to clinical care could have a substantial impact on cost and health at the population level, but are not well powered for discovery on a small scale. If improved reference intervals lead to a more precise diagnosis of hypertension and ultimately reduce the number of individuals treated by 0.1% per year, this represents a cumulative effect of 65,000 individuals (*61*), but would not be detectable in a trial enrolling 1000 people. We did not examine the cost effects of such changes in reclassifying tests and patients. This examination should be done in a more systematic work, which will also include more than the three associated outcomes tested here.

## Materials and Methods

### Data extraction and cleaning

UCSF uses the Epic EHR system, which was launched in August 2012. De-identified structured laboratory data, diagnosis and procedure codes, encounter data, and demographics were extracted from EHRs for encounters between August 2012 and May 2017. We excluded patients <18 or >72 years. For some measurements, reported values are of the form “>x” or “<x” where ‘x’ is a reference number. We replaced x values by increasing or decreasing them by 10% as applicable.

Data were transformed to the same scale. First, in order to match different units, unit designations were transformed to lower case. Next, measurements were transformed to match the same scale. For example, values reported in cells/uL were transformed to cells x10^9^/L by dividing it by 1000.

Healthy encounters were selected by a list of ICD10 codes representing no illness (see Supplemental Table 1). Encounters associated with any other ICD10 code were excluded from the healthy encounter set. Patients older than 60 were excluded from the healthy set.

In the healthy cohort, a single random encounter was picked for each subject. In the case of multiple measurement for the same lab test in one encounter, the median was taken. For general population encounters (healthy and non-healthy patients), only the first measurement of each lab test was taken for each patient.

Patients with unknown/declined race or ethnicity were (Table 1) were included in the healthy cohort, which was used to define EHR-calibrated reference intervals, but were excluded from SIRE-specific intervals calibration.

### Reference intervals

Outliers were detected as proposed by Dixon (*62*) and Reed et al. (*5*). For each test and sex, we defined *IQR* = *Q*_3_ − *Q*_1_ where *Q*_*i*_ was the *i*^*th*^ quartile of the data. Values bellow or above 3 ∗ *IQR* of *Q*_2_ or *Q*_3_ respectively were considered outliers and removed.

According to current practice, a reference interval is defined by collecting ~120 healthy samples from a population. Values between the 2.5^th^ and 97.5^th^ percentiles become the reference interval. Several tests only specify a lower or upper threshold. In these cases, we defined the reference interval as being below the 5^th^ percentile or above the 95^th^ (e.g., LDL Cholesterol < 130 mg/dL and HDL Cholesterol > 39 mg/dL).

Changes in intervals were defined as:

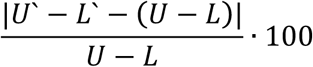

Where U‵, L‵, U, L were EHR-based upper, lower and original upper and lower thresholds.

### Statistical tests

In order to test for statistically significant differences in average values across SIREs, we performed analysis of variance using two tests: one-way ANOVA and the Kruskal-Wallis test (*63*). In the one-way analysis of variance, we added a patient’s SIRE and age to the linear model, as well as the lab measurement as the dependent variable. The Kruskal-Wallis test does not support multiple independent variables, so we first computed residuals of the linear model where lab measurements were dependent variables and age and SIRE were independent. Residuals were then set as dependent variables in the Kruskal-Wallis test with SIRE as the independent one. Multiple test correction was performed using the Benjamini–Hochberg procedure (*64*).

### Outcome regression

We used Cox and logistic regressions to evaluate predictive power and the relationship between a test and a future diagnosis. Logistic regression tests the effect of independent variables with a binary outcome, while Cox regression models censored time to event. Patients already diagnosed with the outcome of interest or diagnosed during the tested encounter were excluded. In each test, we adjusted for SIRE and also performed a statistical test to determine if SIRE was associated with the outcome. To determine if the relationship between lab test value and outcome was different between SIRE groups, we repeated the analyses including an interaction term between the lab test values and the SIRE groups. We tested if regression by SIRE group separately performed better than all samples together. We compared 4 sets of features, given in *R(65)* notation: ‘~’, ‘+’ and ‘*’ indicate dependent variables, independent variables, and interaction of two variables, respectively:

- Y~X+Sex
- Y~X+Age+Sex
- Y~X+Age+Sex+SIRE
- Y~X+Age+Sex+SIRE+SIRE*X

Here, Y denotes outcome event (Table 4). In the logistic regression, Y is binary, while in Cox regression Y is right-censored time to event in days. X is the actual lab value. The interaction term “SIRE*X“ in the 4^th^ set of features models the effect of SIRE on the outcome differently for different values of X.

The assessment of statistical significance in the Cox and logistic regression models was done by likelihood-ratio test of nested models, defined as 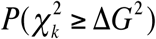 where 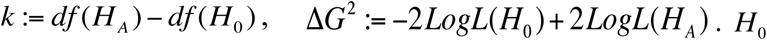 is the reduced model, *H_A_* is the alternative model, and 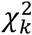 is the chi-squared distribution with 9 degrees of freedom (*df*).

Three lab-specific outcomes were used. Dialysis, as representing kidney failure, for creatinine, T2D for HbA1c, and bacterial infections for WBC. The common bacterial infection used were Pneumonia, Cellulitis, Pyelonephritis or Urinary tract infection (ICD10 codes: J13.*-J18.*, L03.*, N10 or N39.0). Patients having dialysis / T2D record prior to first relevant lab measurement were removed. Kidney failure or T2D diagnoses occurring more than a year from first lab measurement and infection occurring more than 10 days were not considered as positive.

### Cox regression comparison

Non-nested Cox regression models were compared using the Akaike information criterion

(*33*) (AIC). AIC balances the complexity of the model and its predictability. We estimated this parameter by training random sampling 80% of the data and testing using the other 20% samples, and repeated it 1000 times. We reported mean discrimination and standard deviation.

## Supplementary Materials

*Fig. S1. Comparison of new and original reference intervals.*

*Table S1. ICD10 codes*.

List of ICD10 codes used to define healthy patients, and the number of unique encounters and patients associated with each code.

*Table S2. Most common lab tests statistics*.

List of most common lab tests, indicating the number of patients, encounters, measurements for healthy and general statistics, original reference interval and EHR-calibrated reference interval and log ratio of the two intervals.

*Table S3. Complete analysis of variance*.

Most common lab test differences across SIRE groups. Summary statistics and analysis of variance results.

*Table S4. Confusion matrices*.

The number of true/miss classified samples for logistic regression with four sets of features when tested for specific outcomes. For both lab tests, prediction performance improves for each added feature (see Methods). Confusion matrix for WBC was not computed due to low power.

*Table S5. SIRE-specific reference intervals*.

SIRE-specific reference intervals and the fraction of low/normal/high measurements.

## Acknowledgments

We would like to thank Alan Wu and Matthew Kan for fruitful discussions, Alan Wu for reviewing the manuscript, Debajyoti Datta and Merav Heshin-Bekenstein for clinic-related advice, and Dana Ludwig and the UCSF Information Technology Services Academic Research Systems team for Clinical Data Research Consultations and data extraction. Research reported in this publication was supported by the National Institute of General Medical Sciences of the National Institutes of Health under award number R01GM079719. The content is solely the responsibility of the authors and does not necessarily represent the official views of the National Institutes of Health.

---

**Fig. S1.**
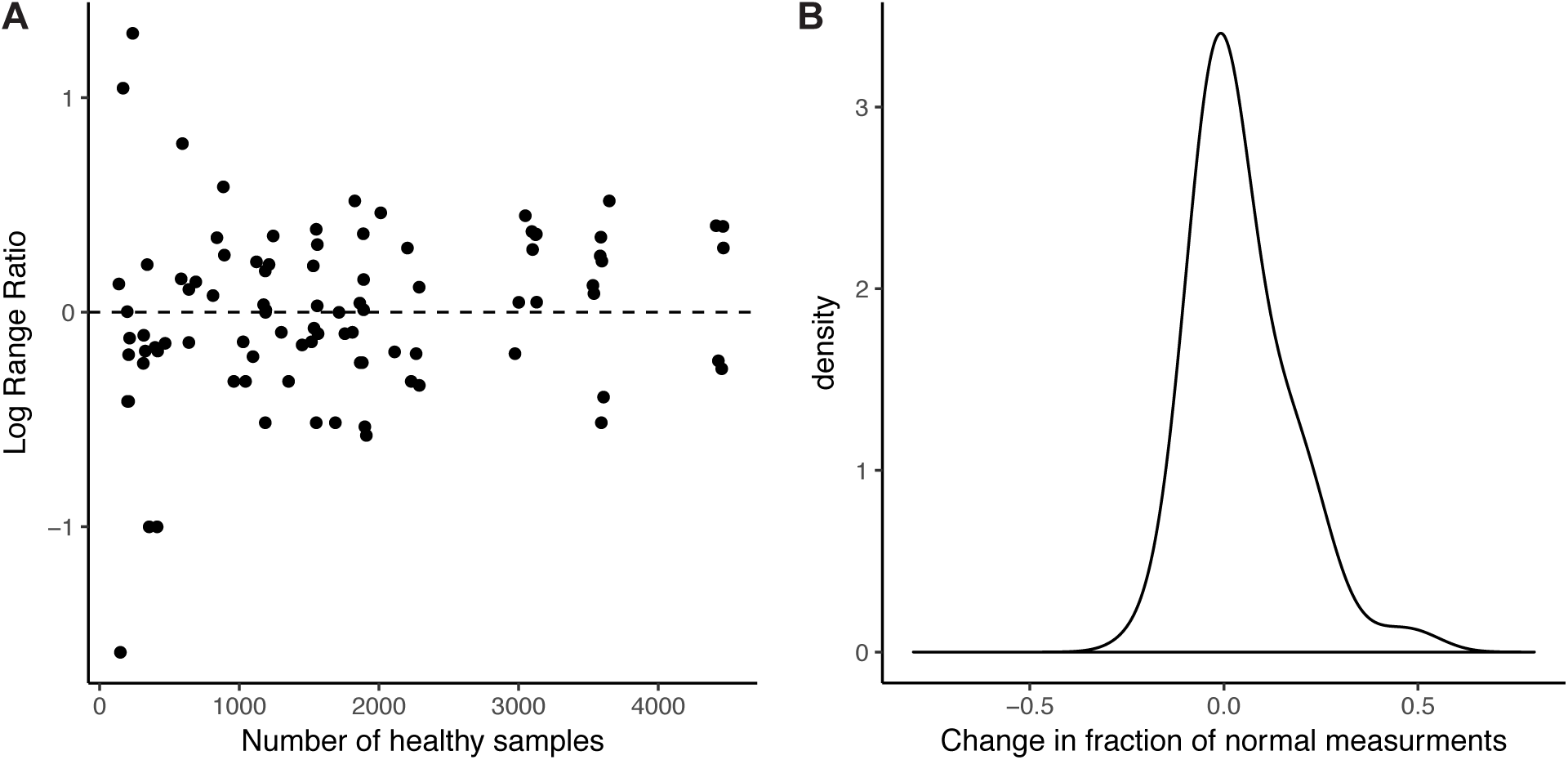
Comparison of new and original reference intervals.

*Table S1. ICD10 codes*.

*Table S2. Most common lab tests statistics*.

*Table S3. Complete analysis of variance*.

*Table S4. Confusion matrices*.

*Table S5. SIRE-specific reference intervals*.

